# Myofiber-specific FoxP1 knockout protects against pancreatic cancer-induced muscle wasting in male but not female mice

**DOI:** 10.1101/2024.09.17.613547

**Authors:** Martin M Schonk, Jeremy B Ducharme, Daria Neyroud, Rachel L Nosacka, Haley O Tucker, Sarah M Judge, Andrew R Judge

## Abstract

Cancer cachexia affects up to 80% of cancer patients and results in reduced quality of life and survival. We previously demonstrated that the transcriptional repressor Forkhead box P1 (FoxP1) is upregulated in skeletal muscle of cachectic mice and people with cancer, and when overexpressed in skeletal muscle is sufficient to induce pathological features characteristic of cachexia. However, the role of myofiber-derived FoxP1 in both normal muscle physiology and cancer-induced muscle wasting remains largely unexplored. To address this gap, we generated a conditional mouse line with myofiber-specific ablation of FoxP1 (FoxP1^SkmKO^) and found that in cancer-free mice, deletion of FoxP1 in skeletal myofibers resulted in increased myofiber size in both males and females, with a significant increase in muscle mass in males. In response to murine KPC pancreatic tumor burden, we found that myofiber-derived FoxP1 is required for cancer-induced muscle wasting and diaphragm muscle weakness in male mice. In summary, our findings identify myofiber-specific FoxP1 as a negative regulator of skeletal muscle with sex-specific differences in the context of cancer.

**NEW & NOTEWORTHY:** Here we identify myofiber-derived FoxP1 as a negative regulator of skeletal muscle with sex-specific effects in cancer. Under cancer-free conditions, FoxP1 knockout increased myofiber size in male and female mice. However, in response to pancreatic cancer, FoxP1 was required for muscle wasting and weakness in males but not females. This highlights the need to consider sexual dimorphism in cancer-induced muscle pathologies and provides evidence suggesting that targeting FoxP1 could help mitigate these effects in males.

## INTRODUCTION

Cancer cachexia is a metabolic condition characterized by progressive skeletal muscle wasting and body weight loss^1^, leading to reduced physical activity and quality of life, poor response to chemotherapy, and increased risk of developing serious comorbidities^2^. Cachexia is also a strong predictor of mortality, and may cause up to 30% of all cancer-related deaths^3^, in part due to respiratory and/or heart failure associated with diaphragm and cardiac muscle atrophy and weakness^4,5^. Despite these devastating consequences, due to the complexity of its etiology there are currently no FDA or EMA-approved treatments for cachexia^6^. One means to improve the likelihood of effective treatments being developed is to continue to improve our understanding of the mechanisms causative in cachexia.

Our laboratory focuses on the skeletal muscle pathology component of cancer cachexia and has previously shown that inhibiting Forkhead box O (FoxO)-dependent transcription prevents muscle atrophy induced by Lewis lung carcinoma (LLC) tumors^7^ and colon-26 adenocarcinoma (C26) tumors^8^. In the C26 model, this inhibition of atrophy was linked to the suppression of well-established atrophy gene networks in skeletal muscle^8^.

When probing into the specific differentially expressed genes in muscle of cachectic C26 mice that were FoxO-dependent, we discovered that the Forkhead box (Fox) family member FoxP1, a ubiquitously expressed transcription factor that has been most widely studied for its role as a transcriptional repressor, is a FoxO target gene in the muscles of C26 mice^8^. FoxP1 has been shown to regulate proliferation and differentiation of various cells types, including immune cells^9^, mesenchymal stem cells^10^, adipocytes^11^ and cardiomyocytes^12,13^. With regards to the latter, FoxP1 has been shown to play a crucial role in cardiac muscle development and physiology, and thus, whole-body *Foxp1* knockout is embryonically lethal in mice due to cardiac defects^13^. In patients with heart failure, FoxP1 protein levels are increased in cardiomyocytes^14^ and FoxP1 mutations have been found in patients with congenital heart defects^15^. However, the physiological importance of FoxP1 in skeletal muscle tissue was largely unknown until recently. Published work from our lab demonstrated that FoxP1 is increased in skeletal muscle of cachectic patients with pancreatic cancer, as well as several mouse models of cancer cachexia, including the C26, LLC, and the human L3.6pl xenograft model^16^. Based on these findings we generated myofiber-specific FoxP1 overexpressing mice which revealed that FoxP1 upregulation in muscle is sufficient to induce severe body and skeletal muscle wasting, as well as myopathy and weakness, causing death in some mice, recapitulating the pathological features and lethality of cancer cachexia^17,18^. Conversely, knockdown of FoxP1 in muscle tissue, using a shRNA driven by an ubiquitous promoter, attenuated C26 tumor-induced muscle wasting. In the current work, we extend these findings by further investigating the requirement of myofiber-derived FoxP1 in a mouse model of pancreatic cancer. To do this, we generated conditional skeletal myofiber-specific FoxP1 knockout (FoxP1^SkmKO^) mice and orthotopically injected them with KPC pancreatic cancer cells to determine the extent to which myofiber-specific FoxP1 is required for pancreatic cancer-induced skeletal muscle wasting and weakness. Importantly, since sex-specific phenotypes are documented in cachectic mice and humans with pancreatic cancer, we conducted studies in both male and female mice. Moreover, we also determined the extent to which myofiber-specific FoxP1 deletion affects muscle fiber size, and muscle mass, in cancer-free male and female mice.

## MATERIALS AND METHODS

### Animals

The University of Florida Institutional Animal Care and Use Committee approved all animal procedures. Animals were provided with ad libitum access to food and water and housed in a temperature-controlled and humidity-controlled facility on a 12h of light/dark cycle. Myofiber-specific FoxP1 knockout (FoxP1^SkmKO^) mice were generated by crossing FoxP1 floxed mice (FoxP1^f/f^), which have been described extensively elesewhere^19–22^, with ACTA-Cre transgenic mice (purchased from the Jackson Laboratory, #006149) which express Cre recombinase under the control of the human alpha-skeletal actin promoter. To note, the FoxP1^f/f^ mice contain a YFP transgene preceded by a floxed stop codon, making YFP expression a reliable reporter of successful recombination. FoxP1^f/f^ mice were used as FoxP1-expressing controls and referred as wild-type (WT) hereafter.

### Model of pancreatic cancer cachexia

Pancreatic cancer cells (KPC FC1245) isolated from the tumor of a LSL-Kras^G12D/+^; LSL-Trp53^R172H/+^; Pdx-1-Cre mouse were obtained from Dr. David Tuveson (Cold Spring Harbor Laboratory, NY, USA)^23^. KPC cells were cultured in Dulbecco’s Modified Eagle Medium supplemented with 10% fetal bovine serum, 1% penicillin and 1% streptomycin at 37 °C in a 5% CO_2_ humidified incubator. 10-week old male, as well as 17-23-week old and 1-year old female WT and FoxP1^SkmKO^ mice were orthotopically injected into the pancreas with 2.5 × 10^5^ KPC cells diluted into 50 µl of sterile saline solution (KPC groups) or saline alone (Control groups), as described previously^24^. Male mice were euthanized 12 days (for muscle function) or 14 days after injections for tissue harvest, and female mice were euthanized 13 days after injections. We have previously shown that KPC tumor bearing mice injected with this cell number, in our hands, develop cachexia 12 days post-injection and reach IACUC-mandated endpoint 15-17 days post-injection^24^.

### Ex vivo muscle function assessment

Ex vivo muscle function was assessed at the Physiological Assessment Core of the University of Florida, as previously described^16^. Briefly, diaphragm muscle strips were mounted on a force transducer placed in a gas-equilibrated (95% O2;5% CO2) 22°C bath of Ringers solution, attached to a dual mode force transducer (Aurora Scientific). After determination of muscle optimal length, maximum isometric tetanic force was determined through a 500-ms stimulation train at 150 Hz. After a 5 min of rest period, diaphragm strips were stimulated at increasing frequencies to generate a force-frequency relationship.

### Histology

Skeletal muscle tissues were embedded in optimal cutting temperature compound (OCT), frozen in liquid nitrogen-cooled isopentane and stored at −80°C until use. Skeletal muscles were cut across the midbelly and sectioned at −20 °C using a Microm HM 550 Cryostat (Microm International) to obtain 10μm cross-sections onto microscope slides, that were either immediately stained with haematoxylin & eosin (as described previously^25^) or stored at −80°C. For Picrosirius Red staining, sections were fixed in Bouin’s fluid at 37°C for 30min then incubated in Picrosirius red solution for 90 minutes. Subsequently, sections were rinsed sequentially in acidified water, Picric Alcohol, and ethanol solutions of varying concentrations, before clearing in xylene and mounting with Cytoseal XYL mounting media. Wheat germ agglutinin (WGA) antibody conjugated with Alexa Fluor 594 (Invitrogen W11262) was used to identify myofibers for measurement of muscle fiber cross-sectional area (CSA) as described previously^16^. Slides were imaged using a Leica DM5000B microscope (Leica Microsystems).

To determine CSA, WGA-stained muscle images were processed using cellpose and the ‘LabelsToRois’ ImageJ plug-in^26^. ImageJ ROI color coder was used to generate representative images where myofibers are colored on a gradient based on their CSA. The percentage of muscle area positive for WGA staining was used to examine differences in extracellular matrix.

### RNA isolation and RT-qPCR

RNA isolation was performed by bead homogenization (BEAD MILL4, Fisher Scientific) of whole TA muscle in TRizol (Invitrogen) followed by extraction with chloroform and precipitation with isopropanol. RNA samples were treated with DNase I then reverse-transcribed to generate cDNA using the SuperScript IV Reverse Transcriptase kit (Thermofisher). TaqMan primers (Applied Biosystems) were used to amplify cDNA and detect expression levels of *FoxP1* (Mm00474848_m1) and *YFP* (Mr04097229_mr). qPCR was done on QuantStudio3 (Applied Biosystems) and quantification was performed using the standard curve or ΔΔCt method.

### Statistical analyses

Data normality was tested by the Shapiro-Wilk test to determine the use of parametric or non-parametric statistical tests. Two-tailed unpaired t-tests or Mann– Whitney U tests were performed accordingly for comparing two groups. Significance level was set at 0.05. CSA data were binned and fit with a Gaussian least squares regression and significance was determined by calculating the extra sum-of-squares F test. Two-way mixed ANOVA was used to test for differences in forces produced at increasing frequencies. Data are presented as mean ± standard error. Statistical analyses and generation of figures were performed using Prism 10 software (GraphPad Software).

## RESULTS

### Confirmation of myofiber-specific FoxP1 knockout

To test for successful Cre-mediated recombination in muscle we measured *YFP* mRNA, which was induced in FoxP1^SkmKO^ mice, but not in FoxP1^f/f^ mice (Figure 1A). *FoxP1* mRNA levels were significantly lower in the muscles of FoxP1^SkmKO^ mice compared to WT mice (Figure 1B), but still detectable. This residual FoxP1 likely derives from non-myofiber cells since *FoxP1* mRNA is highly expressed in all cell types identified in mouse TA muscle (Figure 1C – data extracted from Myoatlas^27^). Together, these data support deletion of myofiber-specific FoxP1 in FoxP1^SkmKO^ mice used in this study.

**Figure 1:**
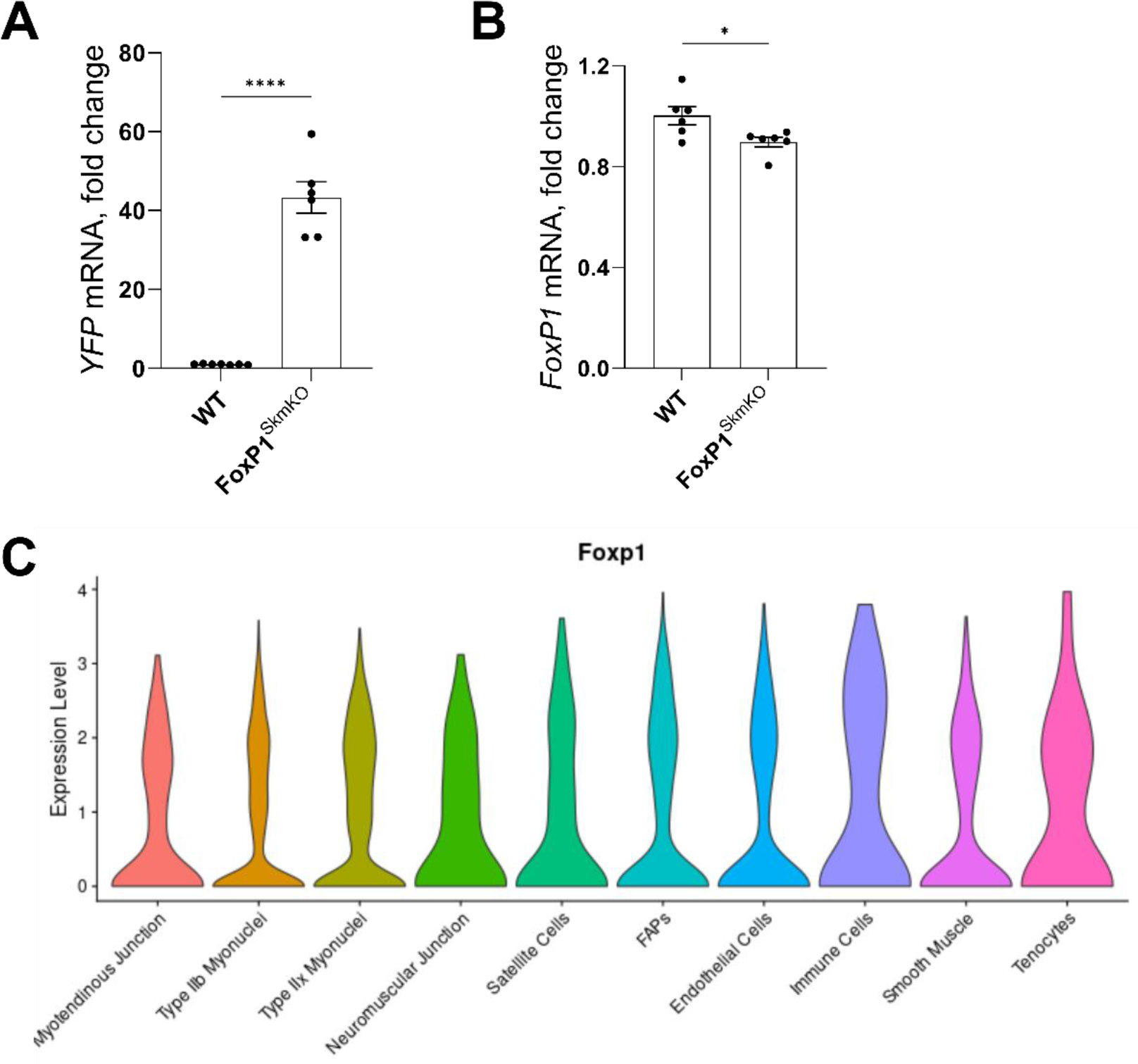
Myofiber-specific FoxP1 knockout. *YFP* (A) and *FoxP1* (B) mRNA levels in mouse TA muscle from WT and FoxP1^SkmKO^ mice. Difference between groups were tested using two-tailed unpaired t-tests (* p<0.05; **** p<0.0001). Data are reported as mean ± SEM. N=5-6 mice/group. C) Single-nucleus RNA-sequencing data in TA muscle of 5-month extracted from the Myoatlas^27^ demonstrate that *FoxP1* mRNA is expressed in cell types identified.

### Muscle-specific FoxP1 knockout increases muscle fiber size in male and female mice

To determine the effect of FoxP1 deletion in skeletal myofibers on muscle mass, we harvested tissue samples from 10-week old FoxP1^SkmKO^ and WT male and female mice (Figure 2A-D). Both TA and *soleus* (SOL) muscle mass were significantly larger in male FoxP1^SkmKO^ mice compared to WT mice (TA: 48.2 ± 1.2mg vs. 44.5 ± 0.9mg, SOL 10.7 ± 0.4mg vs. 8.6 ± 0.3mg). While TA (p=0.152) and SOL (p=0.1143) were numerically larger in FoxP1^SkmKO^ compared to WT in female mice, they did not reach statistical significance (TA: 39.1 ± 2.0mg vs. 35.3 ± 1.3mg, SOL 9.1 ± 1.0mg vs. 6.9 ± 0.5mg). There were no differences between genotypes in heart, spleen, and gonadal adipose tissue masses in either sex (Supplementary Figure 1). Myofiber CSA analysis of TA and SOL muscles from FoxP1^SkmKO^ male and female mice showed a rightward shift in their fiber size distribution curves, indicating a greater number of larger fibers compared to their WT counterparts (Figure 2B-D). No difference in fiber count was observed between genotypes (Supplementary Table 1). To further explore whether FoxP1^SkmKO^ mice show any evidence of altered muscle morphology, extracellular matrix, or collagen content we stained the diaphragm, TA, and SOL muscles using H&E, WGA, and Picrosirius Red, respectively, which revealed no differences between genotypes (Supplementary Figure 2). Overall, these data show that life-long skeletal myofiber-specific FoxP1 deletion increases myofiber size in both male and female mice, resulting in a statistically significant increase in muscle mass in male mice, without affecting the mass of other tissues measured.

**Figure 2:**
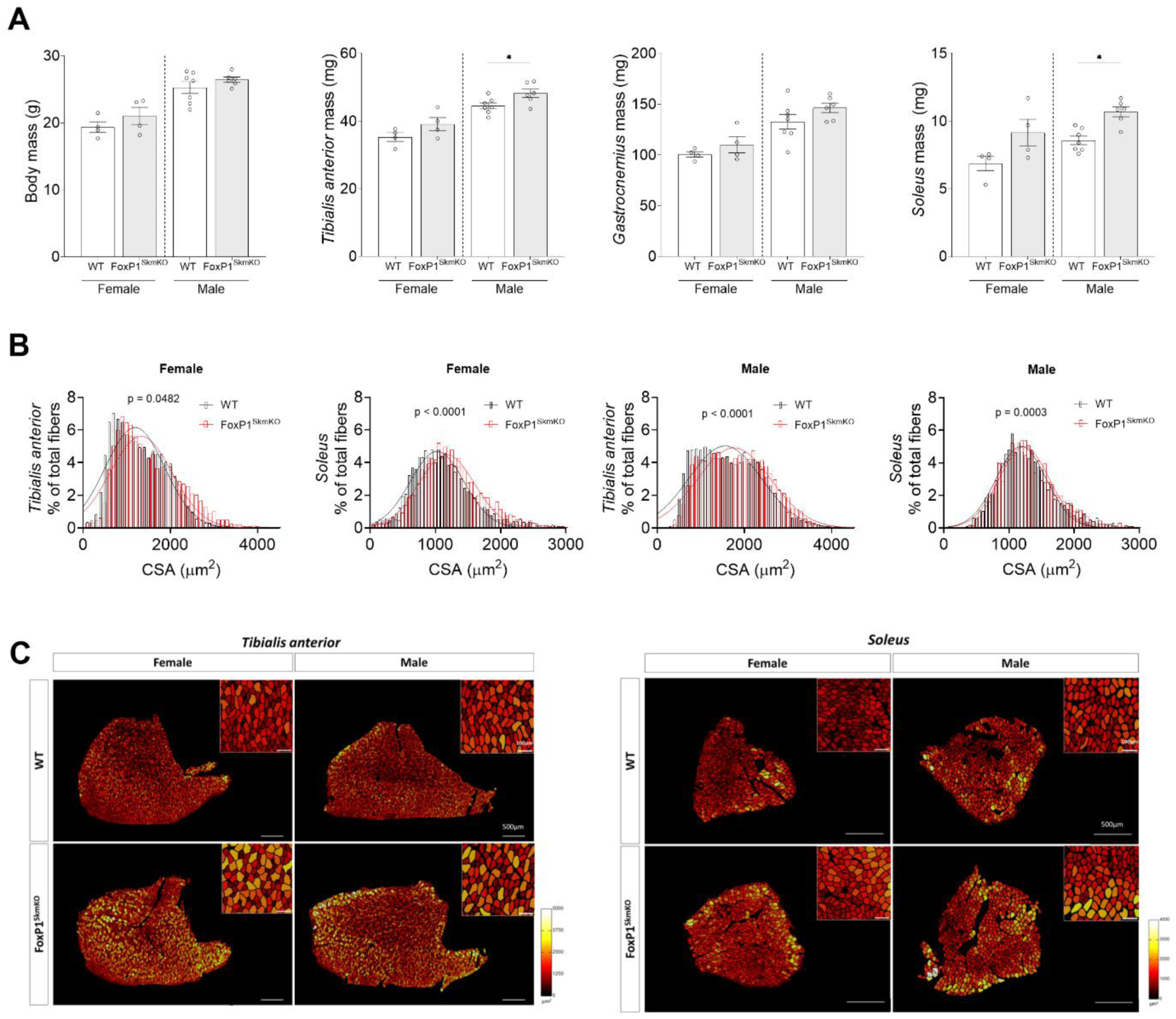
Myofiber-specific FoxP1 knockout increases myofiber size. A) Body and muscle masses from male and female FoxP1^SkmKO^ mice and WT littermates. B) Quantification of TA and soleus muscle fiber cross-sectional area (CSA) reveal significant rightward shift towards larger fibers in FoxP1^SkmKO^ mice. C) Representative images of TA (left) and soleus (right) cross-sections stained for wheat germ agglutinin (WGA), processed for CSA quantification and color-coded based on fiber size. Whole muscle sections (scale bar: 500µm) and zoomed-in inserts (scale bar: 100µm) are displayed. Data are reported as mean ± SEM. Difference between groups were tested using two-tailed unpaired t-tests or Mann–Whitney U tests (* p<0.05). N = 4-7 mice/group.

### Myofiber-specific FoxP1 knockout attenuates cancer-induced muscle atrophy and weakness in male mice

We next aimed to determine the effect of myofiber-specific FoxP1 deletion on pancreatic cancer-induced muscle wasting. To do this, we orthotopically injected male FoxP1^SkmKO^ or WT mice with KPC pancreatic cancer cells, or saline as a control, and harvested tissues 14 days later. There was no difference in tumor mass between genotypes (Figure 3A) indicating that FoxP1 deletion in muscle did not affect tumor growth. Body mass was consistently lower in WT KPC mice, but highly variable in FoxP1^SkmKO^ KPC mice (Figure 3B). Mass of the TA and SOL were significantly decreased in WT KPC mice compared to controls, which was attenuated in male FoxP1^SkmKO^ mice (Figure 3C-E). Consistent with the reduced muscle mass observed in male WT KPC mice vs. control, measurement of TA myofiber CSA revealed a leftward shiftin fiber size distribution, indicating a greater number of smaller fibers in WT KPC mice (Figure 3F, H). No such shift was observed in male FoxP1^SkmKO^ mice (Figure 4G-H), implicating the requirement of myofiber FoxP1 for KPC-induced muscle fiber atrophy in males. Given the protection observed in FoxP1^SkmKO^ KPC mice and our previous findings that FoxP1 overexpression is sufficient to cause muscle weakness^16^, we also tested whether FoxP1 is required for cancer-induced muscle weakness of the diaphragm. Maximum specific tetanic force was decreased by 14.2%, and the force–frequency relationship was lower in WT KPC mice compared to WT control mice (Figure 3I-J). This muscle weakness induced by KPC tumors was prevented in FoxP1^SkmKO^ mice (Figure 3I,K).

**Figure 3:**
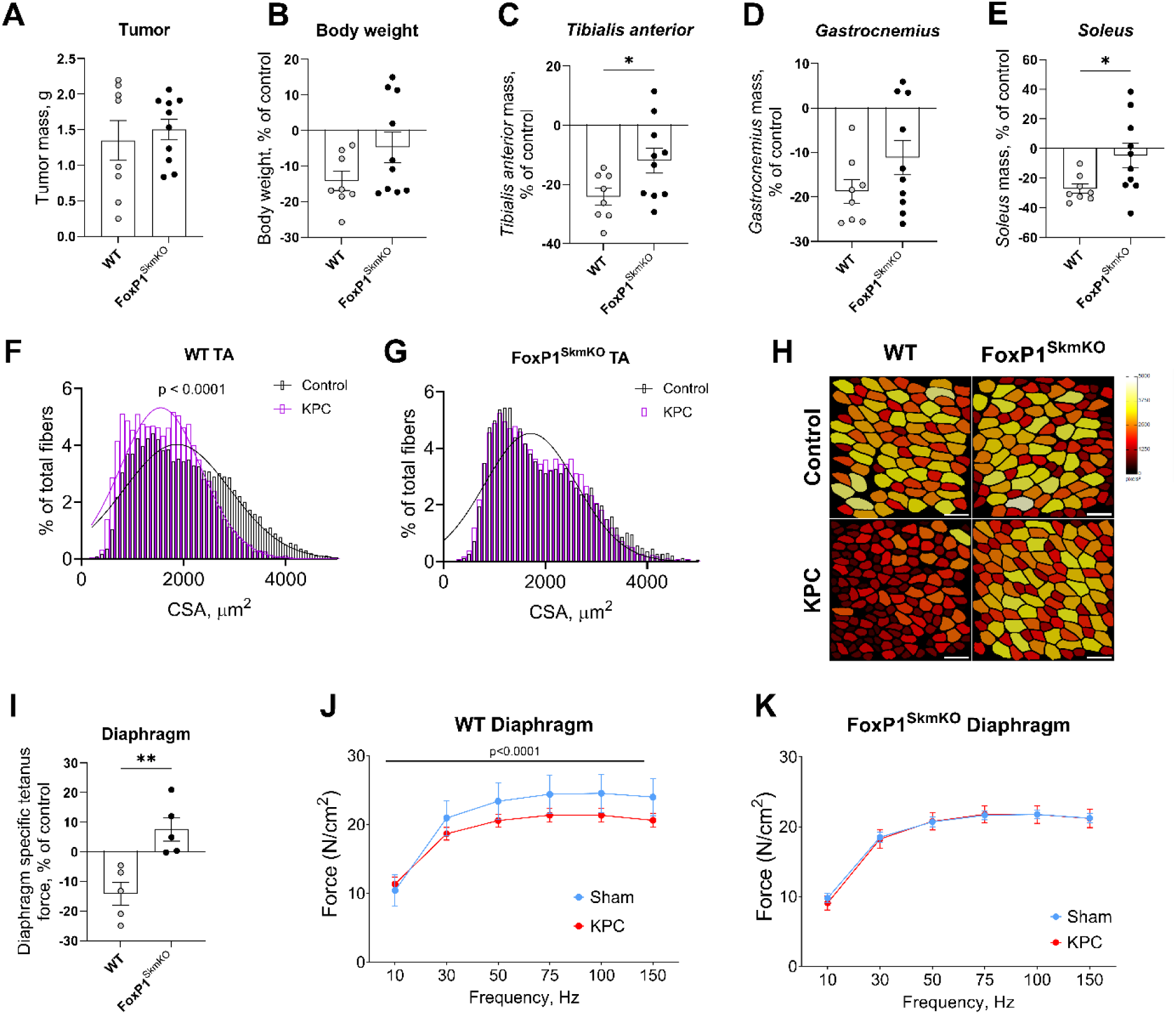
Myofiber-specific FoxP1 deletion protects against pancreatic cancer-induced muscle atrophy and weakness in male mice. A) Tumor weight from WT and FoxP1^SkmKO^ mice at experimental endpoint. B-E) Change in body and skeletal muscle weights of KPC tumor bearing WT and FoxP1^SkmKO^ mice relative to their respective cancer-free genetic controls. F-G) Quantification of TA muscle fiber cross-sectional area (CSA) in WT KPC compared to WT controls (F) and FoxP1^SkmKO^ KPC mice compared to FoxP1^SkmKO^ controls (G). CSA data binned and fit with a Gaussian least squares regression. Significance was determined by calculating the extra sum-of-squares F test. H) Representative images of muscle sections, with fibers color-coded based on CSA. Scale bar: 100µm. I-K) Ex vivo muscle function assessment of diaphragms strips from WT and FoxP1^SkmKO^ mice injected with KPC cells or saline. Data displayed are specific tetanic force (I) and force–frequency relationships (J-K). Two-way mixed ANOVA was used to test for differences in forces produced at increasing frequencies. Data are reported as mean ± SEM. * p<0.05; ** p<0.01. N = 4-10 mice/group.

**Figure 4:**
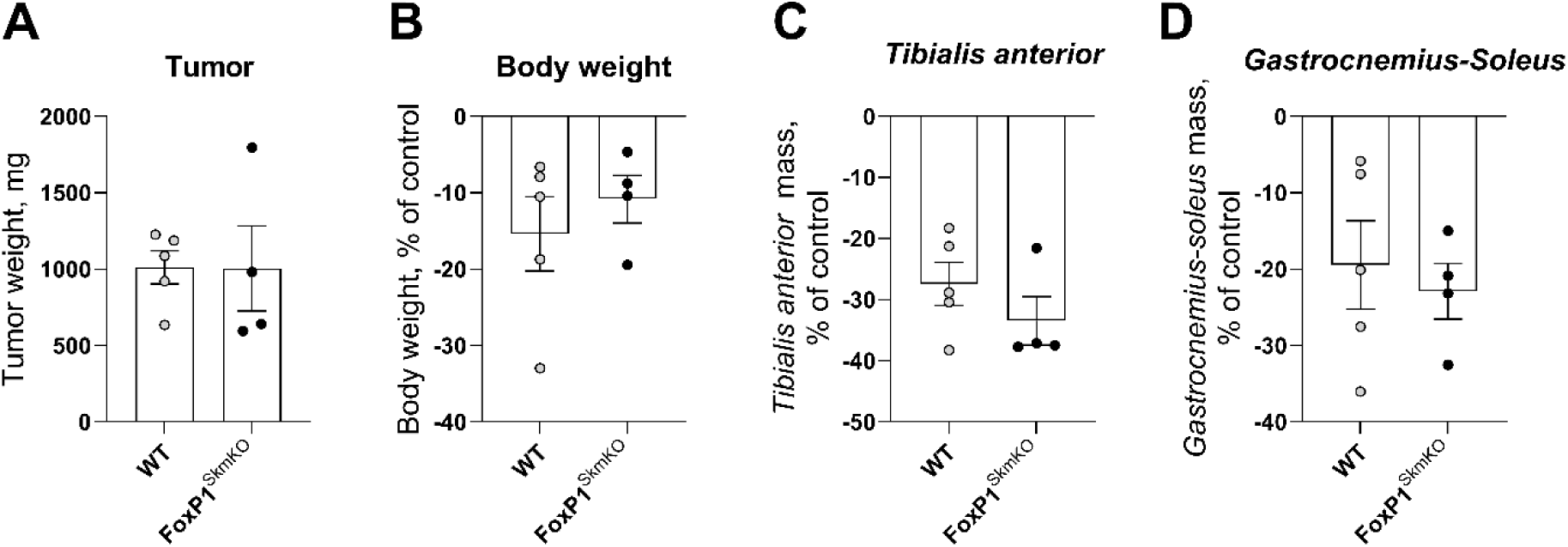
Myofiber-specific FoxP1 deletion does not protect against pancreatic cancer-induced muscle atrophy in female adult mice. A) Tumor mass was no different between genotypes. Change in body mass (B) and skeletal muscle masses (C-D) of KPC tumor bearing WT and FoxP1^SkmKO^ mice normalized to their cancer-free genotype controls. Difference between groups were tested using two-tailed unpaired t-tests. N = 4-5 mice/group. Data are reported as mean ± SEM. All mice were aged 17-23 weeks.

### Myofiber-specific FoxP1 knockout does not protect against cancer-induced muscle atrophy in female mice

Lastly, to determine the extent to which any sexual dimorphism exists in the requirement of FoxP1 for cancer-induced muscle wasting, we injected adult (20-week old) female FoxP1^SkmKO^ and WT mice with KPC cells and evaluated tissue mass. Tumor size was no different between genotypes (Figure 4A), and body and tissue masses were similarly decreased in WT KPC and FoxP1^SkmKO^ KPC mice (Figure 4B-D). Since these results diverged from our findings in male mice and suggested that FoxP1 is not required for cancer-induced muscle atrophy in female mice, we repeated the experiment in female mice, this time using slightly older mice (1-year old). These experiments confirmed our original findings, with comparable tissue wasting in WT and FoxP1^SkmKO^ KPC mice (Supplementary Figure 3).

## DISCUSSION

The present study aimed to investigate the requirement of myofiber-specific FoxP1 in regulating skeletal muscle mass in male and female mice under both normal control conditions and in response to pancreatic cancer. This aim was based, at least in part, on previous findings from our lab which showed that FoxP1 levels are increased in the muscles of cachectic mice and people with pancreatic cancer, and that myofiber-specific FoxP1 overexpression is sufficient to cause muscle wasting, myopathy, and muscle weakness in normal, control conditions^16^. The current study extends these findings by demonstrating that life-long myofiber-specific FoxP1 deletion in normal control mice increases muscle fiber size, with similar effects observed in males and females, suggesting that the role of FoxP1 in skeletal muscle of normal control mice may be independent of biological sex. Taken together, these findings identify myofiber-derived FoxP1 as a negative regulator of skeletal muscle fiber size, as evidenced by the reduction in muscle fiber size with FoxP1 overexpression and the increase in muscle fiber size with FoxP1 deletion.

In the context of cancer, we recently demonstrated that knockdown of FoxP1, via intramuscular injection of an AAV9-U6-FoxP1-shRNA vector, attenuates muscle wasting induced by C26 tumors in male mice^16^. However, the U6 promoter driving the shRNA is not cell type-specific and therefore could knockdown FoxP1 levels in both myofibers and other cell types within skeletal muscle. Moreover, our previous finding that FoxP1 levels are increased in cachectic cancer patients derive from pancreatic cancer patients. Herein we demonstrate the requirement of *myofiber-derived* FoxP1 for pancreatic cancer-induced muscle fiber atrophy in male mice. Interestingly, however, when exploring sex as a biological variable we found that myofiber-specific FoxP1 deletion was not protective against KPC-induced muscle wasting in female mice. Interestingly, similar sex-specific differences were recently identified with the Activin pathway in KPC mice, with administration of the engineered soluble Activin receptor, ACVR2B/Fc, attenuating cachexia in male, but not female, mice^28^. Moreover, when exploring the molecular phenotype of muscle from early-stage male vs. female KPC mice, Zhong and colleagues found only 2.6% overlap in differentially expressed genes (DEGs)^28^. Although the authors found greater overlap in DEGs in late-stage cachexia, it is these early changes that likely drive the subsequent pathologies such as muscle wasting and weakness observed at later stage. Thus, the biological processes which drive KPC-induced muscle wasting appear to be different between sexes, which may therefore explain our divergent findings with myofiber-specific FoxP1 knockout in male versus female KPC mice. Future studies investigating myofiber-specific FoxP1 target genes in male vs. female KPC mice are therefore warranted.

Since a primary goal of preserving muscle and myofiber size is to maintain muscle function, we further extended our studies in male mice to measure muscle force production. We performed these experiments in the diaphragm – a muscle essential to respiratory function and survival, and which we have previously shown exhibits significant weakness in mice bearing both KPC and C26 tumors^24,29,30^. Our data show that myofiber-specific FoxP1 knockout prevented the KPC-induced decrease in maximal tetanic force and the downward shift in the force-frequency relationship measured in WT mice, demonstrating a key role of myofiber-specific FoxP1 in KPC-induced muscle weakness. These findings also align with our recent results which showed that myofiber-specific FoxP1 overexpression is sufficient to induce skeletal muscle weakness in normal control mice^16^.

In summary, these data provide support for myofiber-derived FoxP1 as a negative regulator of skeletal muscle fiber size during normal control conditions. Further, our results suggest that FoxP1 expression in myofibers of male, but not female, mice mediates KPC-induced muscle fiber atrophy. These findings provide pre-clinical evidence of sex-specific differences on the role of FoxP1 in myofibers of mice bearing pancreatic tumors, and suggest that the modulation of FoxP1 could be an effective strategy to prevent pancreatic cancer-induced muscle atrophy and weakness in males. Based on these findings, and those of others, future studies should continue to explore sexual dimorphism in cancer-induced muscle pathologies, in both pre-clinical and clinical samples.

## DATA AVAILABILITY

Data will be made available upon reasonable request.

## AUTHOR CONTRIBUTIONS

MMS, DN, RLN, SMJ, and ARJ conceived and designed the research; MMS, JBD, DN, RLN, SMJ, and ARJ performed experiments; MMS, SMJ and ARJ analyzed data, interpreted results of experiments, prepared figures, and drafted the manuscript. All authors edited, revised, and approved the final version of the manuscript.

## REFERENCES

1. Fearon K, Strasser F, Anker SD, et al. Definition and classification of cancer cachexia: an international consensus. Lancet Oncol. 2011;12(5):489–495. doi:10.1016/S1470-2045(10)70218-7

2. Dahele M, Skipworth RJE, Wall L, Voss A, Preston T, Fearon KCH. Objective physical activity and self-reported quality of life in patients receiving palliative chemotherapy. J Pain Symptom Manage. 2007;33(6):676–685. doi:10.1016/j.jpainsymman.2006.09.024

3. Fearon KCH. Cancer cachexia: developing multimodal therapy for a multidimensional problem. Eur J Cancer Oxf Engl 1990. 2008;44(8):1124–1132. doi:10.1016/j.ejca.2008.02.033

4. Nichols L, Saunders R, Knollmann FD. Causes of Death of Patients With Lung Cancer. Arch Pathol Lab Med. 2012;136(12):1552–1557. doi:10.5858/arpa.2011-0521-OA

5. Ferrara M, Samaden M, Ruggieri E, Vénéreau E. Cancer cachexia as a multiorgan failure: Reconstruction of the crime scene. Front Cell Dev Biol. 2022;10:960341. doi:10.3389/fcell.2022.960341

6. Fearon K, Arends J, Baracos V. Understanding the mechanisms and treatment options in cancer cachexia. Nat Rev Clin Oncol. 2013;10(2):90–99. doi:10.1038/nrclinonc.2012.209

7. Reed SA, Sandesara PB, Senf SM, Judge AR. Inhibition of FoxO transcriptional activity prevents muscle fiber atrophy during cachexia and induces hypertrophy. FASEB J Off Publ Fed Am Soc Exp Biol. 2012;26(3):987–1000. doi:10.1096/fj.11-189977

8. Judge SM, Wu CL, Beharry AW, et al. Genome-wide identification of FoxO-dependent gene networks in skeletal muscle during C26 cancer cachexia. BMC Cancer. 2014;14:997. doi:10.1186/1471-2407-14-997

9. Kaminskiy Y, Kuznetsova V, Kudriaeva A, Zmievskaya E, Bulatov E. Neglected, yet significant role of FOXP1 in T-cell quiescence, differentiation and exhaustion. Front Immunol. 2022;13:971045. doi:10.3389/fimmu.2022.971045

10. Li H, Liu P, Xu S, et al. FOXP1 controls mesenchymal stem cell commitment and senescence during skeletal aging. J Clin Invest. 127(4):1241–1253. doi:10.1172/JCI89511

11. Liu P, Huang S, Ling S, et al. Foxp1 controls brown/beige adipocyte differentiation and thermogenesis through regulating β3-AR desensitization. Nat Commun. 2019;10(1):5070. doi:10.1038/s41467-019-12988-8

12. Steiman S, Miyake T, McDermott JC. FoxP1 Represses MEF2A in Striated Muscle. Mol Cell Biol. 2024;44(2):57–71. doi:10.1080/10985549.2024.2323959

13. Wang B, Weidenfeld J, Lu MM, et al. Foxp1 regulates cardiac outflow tract, endocardial cushion morphogenesis and myocyte proliferation and maturation. Dev Camb Engl. 2004;131(18):4477–4487. doi:10.1242/dev.01287

14. Hannenhalli S, Putt ME, Gilmore JM, et al. Transcriptional genomics associates FOX transcription factors with human heart failure. Circulation. 2006;114(12):1269–1276. doi:10.1161/CIRCULATIONAHA.106.632430

15. Chang SW, Mislankar M, Misra C, et al. Genetic abnormalities in FOXP1 are associated with congenital heart defects. Hum Mutat. 2013;34(9):1226–1230. doi:10.1002/humu.22366

16. Neyroud D, Nosacka RL, Callaway CS, et al. FoxP1 is a transcriptional repressor associated with cancer cachexia that induces skeletal muscle wasting and weakness. J Cachexia Sarcopenia Muscle. 2021;12(2):421–442. doi:10.1002/jcsm.12666

17. Porporato PE. Understanding cachexia as a cancer metabolism syndrome. Oncogenesis. 2016;5(2):e200–e200. doi:10.1038/oncsis.2016.3

18. Martin A, Freyssenet D. Phenotypic features of cancer cachexia-related loss of skeletal muscle mass and function: lessons from human and animal studies. J Cachexia Sarcopenia Muscle. 2021;12(2):252–273. doi:10.1002/jcsm.12678

19. Fu NY, Pal B, Chen Y, et al. Foxp1 Is Indispensable for Ductal Morphogenesis and Controls the Exit of Mammary Stem Cells from Quiescence. Dev Cell. 2018;47(5):629–644.e8. doi:10.1016/j.devcel.2018.10.001

20. Feng X, Ippolito GC, Tian L, et al. Foxp1 is an essential transcriptional regulator for the generation of quiescent naive T cells during thymocyte development. Blood. 2010;115(3):510–518. doi:10.1182/blood-2009-07-232694

21. Li S, Weidenfeld J, Morrisey EE. Transcriptional and DNA Binding Activity of the Foxp1/2/4 Family Is Modulated by Heterotypic and Homotypic Protein Interactions. Mol Cell Biol. 2004;24(2):809–822. doi:10.1128/MCB.24.2.809-822.2004

22. Mazet F, Yu JK, Liberles DA, Holland LZ, Shimeld SM. Phylogenetic relationships of the Fox (Forkhead) gene family in the Bilateria. Gene. 2003;316:79–89. doi:10.1016/s0378-1119(03)00741-8

23. Hingorani SR, Wang L, Multani AS, et al. Trp53R172H and KrasG12D cooperate to promote chromosomal instability and widely metastatic pancreatic ductal adenocarcinoma in mice. Cancer Cell. 2005;7(5):469–483. doi:10.1016/j.ccr.2005.04.023

24. Judge SM, Deyhle MR, Neyroud D, et al. MEF2c-dependent downregulation of Myocilin mediates cancer-induced muscle wasting and associates with cachexia in cancer patients. Cancer Res. 2020;80(9):1861–1874. doi:10.1158/0008-5472.CAN-19-1558

25. Senf SM, Howard TM, Ahn B, Ferreira LF, Judge AR. Loss of the inducible Hsp70 delays the inflammatory response to skeletal muscle injury and severely impairs muscle regeneration. PloS One. 2013;8(4):e62687. doi:10.1371/journal.pone.0062687

26. Waisman A, Norris AM, Elías Costa M, Kopinke D. Automatic and unbiased segmentation and quantification of myofibers in skeletal muscle. Sci Rep. 2021;11:11793. doi:10.1038/s41598-021-91191-6

27. Petrany MJ, Swoboda CO, Sun C, et al. Single-nucleus RNA-seq identifies transcriptional heterogeneity in multinucleated skeletal myofibers. Nat Commun. 2020;11(1):6374. doi:10.1038/s41467-020-20063-w

28. Zhong X, Narasimhan A, Silverman LM, et al. Sex specificity of pancreatic cancer cachexia phenotypes, mechanisms, and treatment in mice and humans: role of Activin. J Cachexia Sarcopenia Muscle. 2022;13(4):2146–2161. doi:10.1002/jcsm.12998

29. Neyroud D, Laitano O, Daguspta A, et al. Blocking muscle wasting via deletion of the muscle-specific E3 ubiquitin ligase MuRF1 impedes pancreatic tumor growth. Res Sq. Published online February 9, 2023:rs.3.rs-2524562. doi:10.21203/rs.3.rs-2524562/v1

30. Roberts BM, Ahn B, Smuder AJ, et al. Diaphragm and ventilatory dysfunction during cancer cachexia. FASEB J Off Publ Fed Am Soc Exp Biol. 2013;27(7):2600–2610. doi:10.1096/fj.12-222844

