## Supplementary Figures for "Myofiber-specific FoxP1 knockout protects against pancreatic cancer-induced muscle wasting in male but not female mice"

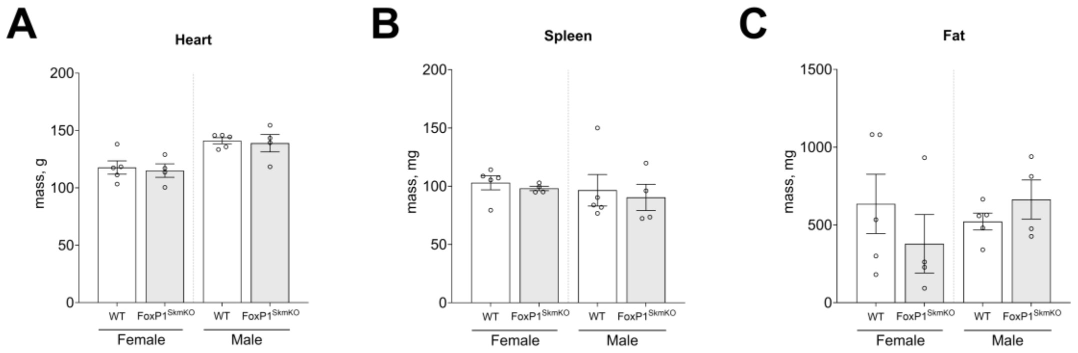


**Supplementary Figure 1: Myofiber-specific FoxP1 knockout does not change A) heart, B) spleen, and C) gonadal adipose tissue mass in FoxP1$^{SkmKO}$ and WT mice.** Difference between groups were tested using two-tailed unpaired t-tests. N = 4-5 mice/group. Data are reported as mean ± SEM.

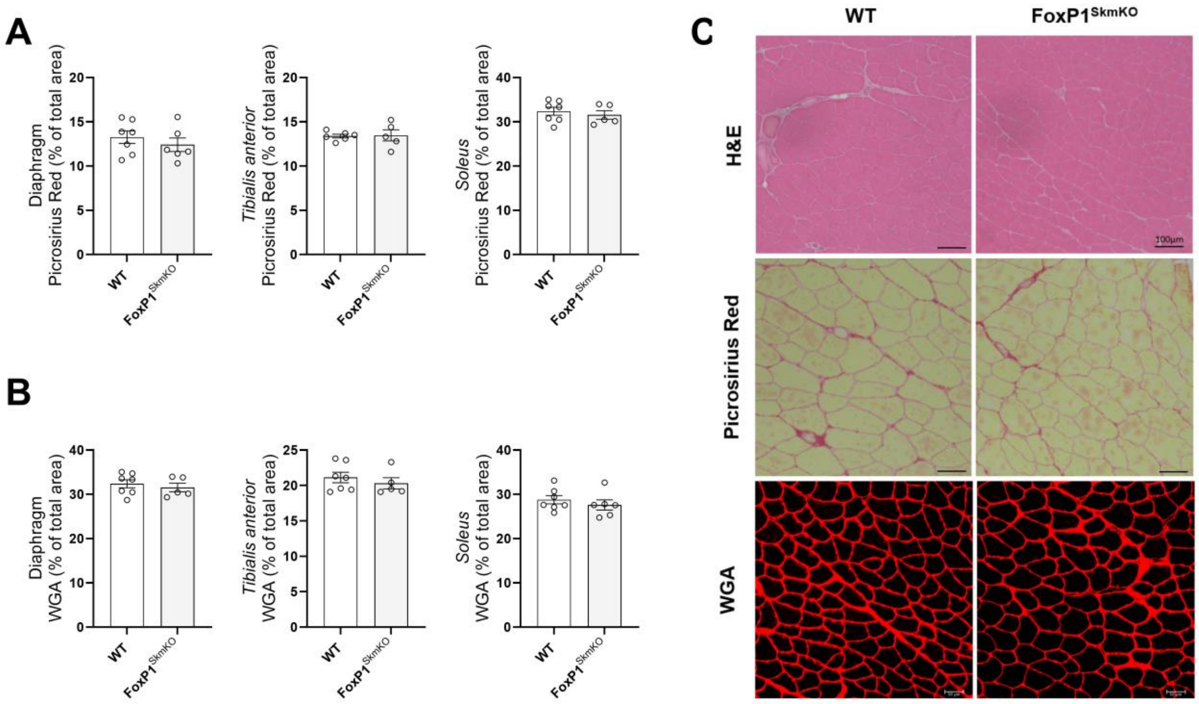


**Supplementary Figure 2: Myofiber-specific knockout of FoxP1 does not alter muscle morphology, extracellular-matrix or collagen content.** Quantification of Picrosirius Red staining (A) and area positive for Wheat germ agglutinin (WGA) staining (B) in Diaphragm, TA and SOL muscles, expressed as % of total area. C) Representative images of Hematoxylin and eosin (H&E), WGA and Picrosirius Red staining of TA muscle cross section (scale bar=100 µm). Difference between groups were tested using two-tailed unpaired t-tests. N = 5-7 mice/group. Data are reported as mean ± SEM.

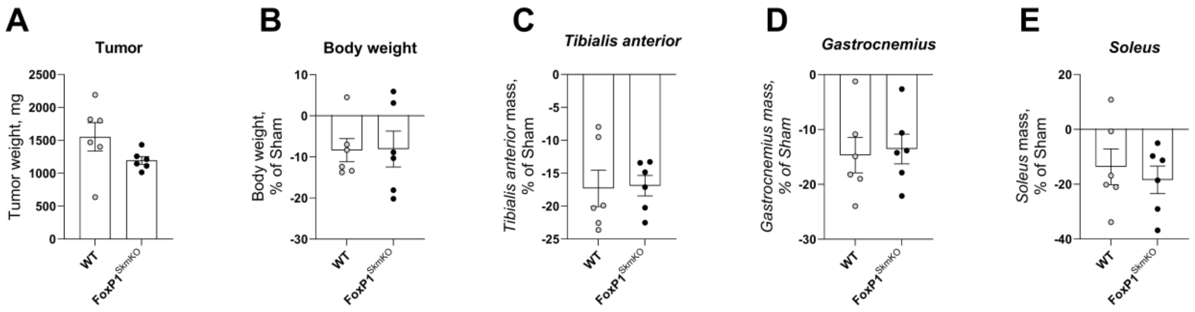


**Supplementary Figure 3: Myofiber-specific FoxP1 deletion does not protect against pancreatic cancer-induced muscle atrophy in female middle-aged mice.** A) Tumor mass was no different between genotypes. Change in body mass (B) and skeletal muscle masses (C-D) of KPC tumor bearing WT and FoxP1$^{SkmKO}$ mice normalized to their cancer-free genotype controls. Difference between groups were tested using two-tailed unpaired t-tests. N = 6 mice/group. Data are reported as mean ± SEM. All mice were 1-year old.

| Sex | Muscle | WT | $FoxP1^{SkmKO}$ | P-value |
|---|---|---|---|---|
| Male | TA | 2343 | 2269 | 0.784 |
| Male | Soleus | 657 | 607 | 0.657 |
| Female | TA | 3020 | 2980 | 0.887 |
| Female | Soleus | 805 | 751 | 0.705 |

**Supplementary Table 1: Average number of fibers of TA and SOL muscles used for cross-sectional area quantification.** Differences between groups were tested with two-tailed unpaired t-tests. N = 4-7 mice/group.
